# European Flint reference sequences complement the maize pan-genome

**DOI:** 10.1101/103747

**Authors:** Sandra Unterseer, Michael A. Seidel, Eva Bauer, Georg Haberer, Frank Hochholdinger, Nina Opitz, Caroline Marcon, Kobi Baruch, Manuel Spannagl, Klaus F.X. Mayer, Chris-Carolin Schön

## Abstract

The genomic diversity of maize is reflected by a large number of SNPs and substantial structural variation. Here, we report the *de novo* assembly of two European Flint maize lines to remedy the scarcity of sequence resources for the Flint pool. EP1 and F7 are important founder lines of European hybrid breeding programs. The lines were sequenced on an Illumina platform at 320X and 225X coverage. Using NRGene´s DeNovoMAGIC 2.0 technology, pseudochromosomes were assembled encompassing a total of 2,463 Mb for EP1 and 2,405 Mb for F7. Structural and functional annotation of the two genomes is currently in progress. The two high-quality *de novo* assemblies complement the existing maize pan-genome and will pave the way for future functional and comparative studies.

## Introduction and organism information

Maize is an important source for food, livestock feed and industrial products. Two of the major germplasm pools exploited in breeding are Dent and Flint with their names referring to different kernel phenotypes [1]. Due to their historic geographical separation and adaptation to different environments, Dent and Flint are genetically divergent with respect to several traits differentiating the two pools, like cold tolerance, early vigour and flowering time. Worldwide, many hybrid breeding programs focus on Dent germplasm, especially the modern high yielding US Corn Belt Dent, whereas breeding programs in cooler regions of Central Europe exploit heterotic effects between Dent and Flint lines. Maize introduced into Europe after the discovery of the New World was substantially influenced by early maturing and cold tolerant North American Flint, but also by Caribbean germplasm [2]. Understanding the genomic differences between germplasm pools may contribute to a better understanding of the complementarity in heterotic patterns exploited in hybrid breeding and of mechanisms involved in the adaptation to different environments.

Maize has a flexible and dynamic genome with a tremendous amount of genetic diversity, which is reflected by a large number of SNPs and substantial structural variation [3-5]. The pan-genome concept presumes that the maize genome comprises genomic segments common to all lines and dispensable segments that can be line-specific or partially shared between lines [6]. High-quality genome sequences are essential for characterizing and exploiting the native genomic variation within maize. To date, a reference sequence of maize exists based on the Dent line B73 [3, 7] as well as a recently published *de novo* assembled genome of the Dent line PH207 [8]. To remedy the scarcity of sequence resources for the Flint pool, two reference sequences were generated *de novo* from inbred lines EP1 and F7. The two Flint lines are important founder lines of European hybrid breeding programs and trace back to the Spanish landrace Lizargarate and the French landrace Lacaune, respectively.

## Genome sequencing information

For Illumina whole genome sequencing, DNA was extracted from leaf material frozen in liquid nitrogen following the protocol of [9]. DNA was extracted from 17 (EP1) or 15 (F7) single plants and then pooled for library construction. The libraries for both samples were prepared following manufacturer´s instructions. We obtained a raw sequencing data coverage of 320X for EP1 and of 225X for F7 based on libraries with five different insert sizes (470 bp, 800 bp, 3000 bp, 6000 bp, and 9000 bp). Contigs, scaffolds and pseudochromosomes were assembled using NRGene´s DeNovoMAGIC 2.0 technology. The resulting 137,249 and 130,426 contigs had an N50 length of 82,295 bp and 96,432 bp for EP1 and F7, respectively. The contig N50 count was 8,811 (EP1) and 7,368 (F7). The N50 length for 71,196 EP1 scaffolds was 6.134 Mb and the N50 length for 77,899 F7 scaffolds was 9.483 Mb. The scaffold N50 counts were 121 and 70, respectively. The final pseudochromosomes encompassed a total of 2,463 Mb (EP1) and 2,405 Mb (F7) including 2,438 Mb and 2,379 Mb ungapped sequence, respectively. Finally, we used BUSCO [10] to assess completeness of the two genome assemblies based on evolutionarily-informed expectations of gene content from 956 near-universal single-copy orthologs. The assemblies of EP1 and F7 revealed 97.3% and 96.1% of complete BUSCOs and 1.6% and 2.4% of fragmented BUSCOs, respectively.

## Conclusions

High-quality *de novo* assembled reference sequences are fundamental for characterizing genomic and functional variation. The two reference sequences presented here will enable the integration of Flint diversity into the maize pan-genome and will pave the way for more detailed structural and functional analyses of Flint germplasm.

## Availability of data

The *de novo* assembled genomes, Zm-EP1-REFERENCE-TUM-1.0 (Zm00010a) and Zm-F7-REFERENCE-TUM-1.0 (Zm00011a), were released in collaboration with the Maize Genetics and Genomics Database (MaizeGDB; http://www.maizegdb.org) and NCBI (BioProjects PRJNA360920 and PRJNA360923). The Whole Genome Shotgun projects have been deposited at GenBank under the accessions MTTA00000000 and MTTB00000000. The versions described here are versions MTTA01000000 and MTTB01000000. Functional annotation supported by RNA-seq data and structural analysis of the two genomes is currently in progress.

The Plant Breeding Group (Technical University of Munich) and the Plant Genome and Systems Biology Group (Helmholtz Center Munich) in collaboration with NCBI and MaizeGDB have released the two maize genomes prior to scientific publication in accordance with guidelines set forth by the Toronto Agreement for prepublication data sharing (Nature. 2009 461:168). The above groups reserve the first right to publish on the available EP1 and F7 data, including but not limited to whole-genome comparisons, genes, structural annotations, functional annotations, and genome-wide association studies, and to improve the sequence and its annotations.

By accessing these data users agree not to publish articles containing whole genome or chromosome scale analyses prior to publication by the Plant Breeding Group (Technical University of Munich) and the Plant Genome and Systems Biology Group (Helmholtz Center Munich). Any redistribution of these data should include the full text of the data use policy.

## Authors’ contributions

CCS, EB, KFXM, MS, GH and FH conceived the study; NO and CM performed sample preparation; KB assembled the genome; MAS conducted data analysis; SU and EB wrote the manuscript. All authors read and approved the final manuscript.

## Funding acknowledgements

This study was funded by the Bavarian State Ministry of the Environment and Consumer Protection (Grant ID TGC01GCUFuE69741; Project BayKlimaFit; http://www.bayklimafit.de/) and the German Federal Ministry of Education and Research (BMBF) within the funding initiative “Plant Breeding Research for the Bioeconomy” (Grant ID 031B0195A; Project MAZE; http://www.europeanmaize.net).

